# Pure yeasts selected from fermenting pear juice from the Lower Austrian Mostviertel region and their suitability for pear wine production

**DOI:** 10.1101/2024.03.05.583456

**Authors:** H. Gangl, G. Tscheik, W. Tiefenbrunner

## Abstract

If spontaneous fermentation is not carried out, commercially available pure yeasts, which are actually intended for the production of wines from grapes, are often used for the production of pear wines. Pear juice is clearly different from grape juice, e.g. due to the significantly greater difference in the ratio of fructose to glucose or the absence of tartaric acid. For this reason, the commercially available pure yeasts may be suboptimal for pear wine production. They were therefore compared with yeasts isolated from pear juice that had been purely cultivated with regard to their suitability for pear wine production.

Overall, the yeasts isolated from fermenting pear juice proved to be more suitable for pear wine production. This is not only the result of the tasting of the wines fermented in 25-litre glass flasks, but also follows very clearly from the multivariate comparison both for the basic chemistry and for the spectrum of aroma components. Density and malic acid concentration were higher in wines produced with yeasts selected from fermenting pear juice. More substances from the "fruity" and especially "fruity-sweet" flavour groups were found in these wines in comparatively higher concentrations.

For pear wine production, it is therefore advisable to isolate suitable yeasts from pear juice for pure cultivation.

## Introduction

The genus *Pyrus* (pear), belonging to the Rosaceae family, probably evolved in the Tertiary period (66 to 55 million years ago) in the mountainous regions of western China (Silva et al. 2014). Phylogenetically, it is close to the genus *Malus* (apple). Distribution and speciation followed the course of the mountains both eastwards and westwards. *P. theobroma* has been described from the Miocene (23 to 5 million years ago) in Austria (Europe), the locality is Parschlug in the district of Bruck an der Mur (Rubtsov 1944).

The domestication of fruits through vegetative propagation began around 6000 years ago, initially in the Caucasus, from where the practice spread to Asia Minor, the Middle East and the Mediterranean. In addition to vines, figs, olives and pomegranates, pears also appear to have been domesticated early on. However, the earliest mention of the pear in the West was made more than 3000 years later by Homer (in China, the cultivation of the pear was mentioned in writing as early as 2000 BC). In the Odyssey, the poet describes the pear as a gift from the gods. Another Greek, Theophrastus (371-287 BC), distinguished wild pears from cultivated varieties. The Romans adopted cultivation, as evidenced in a document written by Portius-Cato (235-150 BC). He distinguishes between six varieties of pears. Terentius Varro (116-27 BC) wrote about the preservation of the fruit. Due to their long storage life, pears were comparatively easy to transport in antiquity and therefore highly sought-after. The Roman Pliny the Elder (23-79 AD) made a significant contribution with his description of more than 40 varieties of pears. He was also the first (as far as is known) to report that wine was made from pears and all varieties of apples.

Cultivation declined at the end of the Roman period, but was increasingly resumed by monasteries and aristocrats from around 600 AD (Normandy initially played a special role) and pears were also used for wine production. Because hygiene was not always excellent, alcoholic drinks were relatively healthy due to the bactericidal effect of ethanol, but the flavour and the relaxing and intoxicating effect certainly played a role in their popularity, as they still do today. Pears have a high nutritional value and are rich in vitamins (A, B1, B2, B3, C) and minerals. *P. nivalis* is particularly important for wine production, *P. communis* for the production of fruit for direct consumption. *P. nivalis* has a higher tannin content. Pear wine has been of great importance in France and Britain for at least 400 years.

Austria has a very large number of indigenous cider pear varieties - many of which were mentioned in writing centuries ago - whose area of origin lies between the Hausruck in Upper Austria and the Traisen in Lower Austria. Carinthia and Styria also have a long tradition of pear wine production.

Since 2014, it has been possible to apply for the "Federal Quality Assurance Number" (“Staatliche Prüfnummer”) for quality fruit wine. Accordingly, it is necessary to have yeasts available for fruit wine that ensure harmonious fermentation. In principle, yeasts from the native "wild strain" of a region are preferable in order to preserve the local characteristics of the fermentation product.

The primary aim of this project was to obtain pure yeasts for pear wine production in order to give fruit wines their characteristic flavours, fullness and typical identity. The question of whether conventional *Saccharomyces* pure culture yeasts, produced for grape wine production, are equally suitable for fruit wines was also to be clarified. Pure culture yeasts enable stable fermentation and can guarantee a constant and very high quality of the end product and are therefore also of interest for pear wine production.

## Method

### Pure cultivation of yeasts

Pear juice from cider pears (Wasserbirne, Dorschbirne, Speckbirne and Grüne Pichlbirne) from the Lower Austrian Mostviertel region was fermented natively in autumn under cellar conditions separately for the individual varieties. Yeast was extracted from various stages of fermentation on two dates (mid-October and early November) for further selection, which was carried out intensively the following year. GYP (glucose, yeast extract, peptone agar) culture medium in Petri dishes (5cm diameter) was used for pure yeast cultivation. The colonies were separated and transferred further until eidonomic-morphological homogeneity was achieved in the cells of a culture. Nine yeasts were finally available.

### Basic analyses of grape juice and wine

To describe the "basic chemistry", the content of sugars (glucose, fructose), acids (titratable and total acid, volatile acid [sum of those acids that can easily "volatilise" in wine, especially acetic acid], citric acid, malic acid and lactic acid; the content of tartaric acid is zero in pear wines) and alcohols (ethanol, glycerol) is given, as well as some physical parameters (sugar gradation, density and pH value). The yeast-available nitrogen (YAN) is important for the fermentation process, the alcohol content increases during fermentation, the sugar content decreases. Density and dry extract content are highly correlated.

The basic parameters were analysed according to Schneyder 1979 or according to the OIV method book.

### Initial substrate

Pear press juice from the previous autumn was used for fermentation. Some basic parameters of the pear juices used for vinification are summarised in Table 1.

**Table 1:** Parameters of the basic analysis carried out for the initial substrate (pear juice). A: first year of vinification, B: second year.

| Parameters | Unit | Pear juice |  |
| --- | --- | --- | --- |
|  |  | A | B |
| Density |  | 1.0674 | 1.0599 |
| Total sugar content | g/l | 133.8 | 123 |
| Fructose | g/l | 99.8 | 96 |
| Glucose | g/l | 34 | 27 |
| Titrateable acid (as malic acid) | g/l | 5.4 | 5.8 |
| Titrateable acid (as tartaric acid) | g/l | 6 | 5.2 |
| Volatile acid | g/l | 0 | 0 |
| Tartaric acid | g/l | 0 | 0 |
| Citric acid | g/l | 1.4 | 2.8 |
| Malic acid | g/l | 4.9 | 4.2 |
| Glycerol | g/l | 0 | 0 |
| pH value |  | 3.63 | 3.6 |
| N yeast available | mg/l | 51 | 11 |

### Control of the fermentation process

In the case of microvinification in the 300 ml conical flask, the fermentation process was controlled by the weight loss of the test sample, from which the total amount of carbon dioxide produced during fermentation can be calculated if the evaporation loss is taken into account. During fermentation in the 100 litre tanks, heat production was used as a measure of fermentation rate.

### Tasting

A tasting of the resulting wines from the second and third year of the study was carried out with a tasting panel of seven tasters. The assessment criterion was marketability. The tasters were not informed of the identity of the yeast used to vinify the pear juice in order to maintain impartiality.

### Aroma analysis

The analysis of the aroma spectrum initially comprised 61 aroma-active substances (in the wines obtained by microvinification in 300ml flasks), which were reduced to 39 after some experience with pear wines. A gas chromatography-mass spectrometry system, quadrupole mass spectrometer from Shimadzu Austria (Korneuburg) GC/MS QP2010, Ultra, was used to analyse the aroma components (aldehydes, esters, alcohols, acids) of the pear wines. A polar capillary column from Zebron (Aschaffenburg, Germany), ZB-WAX plus, length 60m, film thickness 0.50μm, inner diameter 0.25mm, was used as the separation column.

Analyte enrichment was performed by solid phase microextraction (SPME). An 8 ml sample was used and mixed with internal standard (3-decanol). After equilibration (40°C/5 minutes) in the autosampler from CTC Analytics (Zwingen, Switzerland), the adsorption of analyte molecules on the SPME fibre (a 2cm long C/DVB/PDMS fibre carboxene/divinylbenzene/polydimtheylsiloxane, 50μm/30μm) in the gas phase (headspace analysis) was carried out for 30 minutes. This was followed by the actual analysis. The injector temperature was held at 50°C for 3 minutes before being increased by 4°C per minute to 180°C and then to 230°C at a rate of 25°C/min and then held at this temperature for a further 7.5 minutes. The entire cycle lasted 45 minutes. Helium was used as the carrier gas at a constant flow rate of 1.0 ml/min. The result of this procedure is not the concentrations of the flavour components, but their specific ion track (mass fragment) areas (SIM areas).

Principal component analysis (PCA), a statistical method used for dimensional reduction, was applied for the multivariate comparison of the flavours and the basic chemistry of the wines.

### Microvinification (300ml)

Microvinification was carried out in March in 300ml conical flasks at a constant temperature of 20°C with daily recording of the batch weight. Per flask, one of the selected yeast strains was added to 250 ml of mixed pear juice from the region and, for comparison, another flask was added with a commercially available pure cultured grape wine yeast of the species *Saccharomyces cerevisiae* (Pannonia: PAN; K) and one with a yeast isolated from apple juice in the Mostviertel region (GH DT 1 apple A4; C).

Immediately after fermentation, the basic wine parameters were recorded. On the other hand, the remainder was deep-frozen at -36°C for flavour analysis.

Microvinification was primarily used to test the suitability of the selected "pear's own" yeasts, as well as the comparative yeasts, for pear wine production.

### Fermentation in 100-litre tanks without comparative yeasts

One year after microvinification, in September, vinification began in 100-litre fermentation tanks. Only six yeasts were used, which were identical to those used for microvinification; three had been excluded by microvinification as unsuitable for pear wine production. The basic analysis of the starting substrate (pear juice) was carried out previously and is shown in Table 1 (A). This fermentation was carried out without comparative yeasts, i.e. exclusively with yeasts isolated from pear juice. The basic parameters of the wines and an aroma analysis were recorded.

### 25-litre and 100-litre fermentation with comparative yeasts

To supplement the experiments carried out in previous years, pear juice was again fermented, whereby some of the yeasts used were again isolates from native pear juice fermentation. For comparison, however, this time several commercially available pure yeasts were used, which are normally utilised in grape wine production. Fermentation was carried out on 18 wines (nine wines in duplicate) in 25-litre glass flasks at the Haidegg test station, and in two cases in 100-litre tanks in a cellar in the Lower Austrian Mostviertel region, where the vinifications had already taken place the previous year.

Mixed pear juice from the previous year was used as the fermentation medium, consisting of 50% Speckbirne pear must, 35% Grünpichlbirne pear must and 15% Landlbirne pear must. The basic chemistry of the starting substrate is shown in Table 1 B. The test yeasts used were those best evaluated in the previous experiment, isolated from Grüne Pichlbirne, Speckbirne and Wasserbirne as fermentation medium. Yeast **B**, obtained from fermenting Dorschbirne pear juice and judged best in the previous test, was cultivated further in the manner already described and the two results were labelled "Yeast 1" and "Yeast 2". In addition, the following commercially available yeasts were used for comparison: Actiflore Rose, Filtraferm Tropic, Oenoferm Freddo and Oenoferm X-Thiol.

## Results

### Microvinification (300ml)

Microvinification was primarily used to assess whether the yeasts isolated from the spontaneously fermenting pear juice were suitable in principle for the production of pear wines. After pure cultivation, nine strains were finally available. In addition, a yeast **C** isolated from fermenting apple juice and a commercially distributed *Saccharomyces cerevisiae* pure culture yeast Pannonia **K** (EGH 2, Lallemand GmbH) were used for comparison. Table 2 lists all yeasts and also contains the basic assessment.

**Table 2:** Designation of the yeast strains isolated from various spontaneously fermenting pear juices for microvinification. The information on the left serves to reconstruct the selection process; only the individual letters **A**-**K** are of significance for the comparison. A commercially available pure yeast (**K**) and one isolated from fermenting apple juice (**C**) were also used for comparison. The "Quality" column assesses the suitability of the yeast for pear wine production according to the results of the basic analysis presented in Table 3 and the fermentation process shown in Fig. 1.

| Yeast strain |  | Quality | Pear juice |
| --- | --- | --- | --- |
| GH HB B4 DB | B | yes | Dorschbirne |
| GH DT 2 A4 DB | H | no |  |
| GH Zj 2 A3 GPB | F | yes | Grüne Pichlbirne |
| GH REIK 2 A4 GPB | I | yes |  |
| GH Datz B1 GPB | J | yes |  |
| GH Zj 2 B5 SB | E | yes | Speckbirne |
| GH Datz A6 SB | G | no |  |
| GH Zj 2 A WB | D | yes | Wasserbirne |
| GH HB B4 WB | A | no |  |
| GH DT 1 Apple A4 | C | yes |  |
| Pannonia | K | yes |  |

The assessment was based on fermentation efficiency, which can easily be judged by the alcohol content of the finished wine on the one hand and the CO_2_ production during the fermentation process on the other. Although the fermentation of pear juice produces less alcohol than that of grape juice, a content of less than 4% was judged to be insufficient. Three yeasts had to be excluded from further trials, one each isolated from Dorschbirne (**H**), Speckbirne (**G**) and Wasserbirne (**A**).

Table 3 also shows that none of the wines had tartaric acid and that there were very large differences in terms of sugar utilisation, particularly with regard to the ratio of glucose to fructose. For example, yeast **I** (Grüne Pichlbirne) metabolised almost all of the sugar, while others metabolised very little.

**Table 3:** Parameters of the basic analysis that was carried out for the pear wines produced by microvinification (300ml) and mean values for all wines. Fermentation was undertaken with the pure yeasts listed in Table 2.

|  | Wasserbirne |  | Dorschbirne |  | Speckbirne |  | Grüne Pichlbirne |  |  | Apple | Grape |  |
| --- | --- | --- | --- | --- | --- | --- | --- | --- | --- | --- | --- | --- |
| Identity | A | D | B | H | E | G | F | I | J | C | K | mean |
| Density | 1.0419 | 1.0203 | 1.0097 | 1.0533 | 1.0244 | 1.0387 | 1.0149 | 1.0071 | 1.0183 | 1.0212 | 1.017 | 1.024 |
| Alcohol | 1.8 | 4.5 | 5.9 | 0 | 4 | 2.2 | 5.2 | 6 | 4.6 | 4.4 | 4.9 | 3.95 |
| Dried extract | 115.5 | 69.4 | 46.8 | 141.6 | 78 | 109 | 57.7 | 40.4 | 64.4 | 71.5 | 62.4 | 77.88 |
| Fructose | 51.2 | 26 | 7.5 | 68.1 | 34.3 | 47.1 | 17 | 0.8 | 22.6 | 31.3 | 21.5 | 29.76 |
| Glucose | 11.5 | 3.5 | 0.6 | 21.2 | 5.2 | 9.1 | 1.5 | 0 | 1.8 | 3.2 | 2.1 | 5.43 |
| Sugar-free extract | 52.8 | 39.9 | 38.7 | 52.3 | 38.5 | 52.8 | 39.2 | 39.2 | 40 | 37 | 38.8 | 42.65 |
| Extract remainder | 44.3 | 30.9 | 29.4 | 44.3 | 29.8 | 43.8 | 30.2 | 30 | 30.3 | 27.8 | 29.4 | 33.65 |
| Titrateable acid as tartaric acid | 9 | 9.4 | 9.7 | 8.5 | 9.1 | 9.8 | 9.5 | 9.5 | 10 | 9.6 | 9.8 | 9.45 |
| Titrateable acid as malic acid | 8 | 8.4 | 8.7 | 7.6 | 8.1 | 8.8 | 8.5 | 8.5 | 8.9 | 8.6 | 8.8 | 8.45 |
| Volatile acid | 0.4 | 0.3 | 0.3 | 0.4 | 0.3 | 0.6 | 0.4 | 0 | 0 | 0.3 | 0.3 | 0.30 |
| Tartaric acid | 0 | 0 | 0 | 0 | 0 | 0 | 0 | 0 | 0 | 0 | 0 | 0.00 |
| Citric acid | 2.4 | 2.8 | 2.7 | 2.9 | 2.8 | 2.6 | 3.1 | 3.1 | 2.5 | 2.8 | 2.9 | 2.78 |
| Malic acid | 5 | 5.7 | 5.9 | 5.4 | 5.4 | 5.3 | 5.7 | 5.7 | 5.5 | 5.9 | 6 | 5.59 |
| D-lactic acid | 0 | 0 | 0 | 0.1 | 0 | 0 | 0 | 0 | 0 | 0 | 0 | 0.01 |
| L-lactic acid | 0 | 0 | 0 | 0 | 0 | 0 | 0 | 0 | 0 | 0 | 0 | 0.00 |
| Glycerol | 1.4 | 2.5 | 2.8 | 0.8 | 2.8 | 2.1 | 2.7 | 3.1 | 3.7 | 2.5 | 3.1 | 2.50 |
| pH value | 3.5 | 3.42 | 3.48 | 3.47 | 3.43 | 3.39 | 3.44 | 3.52 | 3.45 | 3.42 | 3.43 | 3.45 |
| N yeast available | 10 | 7 | 2 | 0 | 2 | 6 | 0 | 0 | 0 | 8 | 0 | 3.18 |

The fermentation efficiency of the yeasts was also monitored via the CO_2_ loss of the test sample (Fig. 1), which is naturally related to alcohol production via anaerobic fermentation (C_6_H_12_O_6_ **→** 2 C_2_H_5_OH + 2 CO_2_; see e.g. Kinzel 1977). With regard to this loss, it was determined that a production of less than 30g CO_2_ per litre of pear juice must be evaluated as an indication of insufficient fermentation efficiency. Three yeasts are excluded by this criterion, namely **A, G** and **H**, as expected in accordance with the elimination due to insufficient ethanol production. Six yeasts remained for further analyses.

**Fig. 1:**
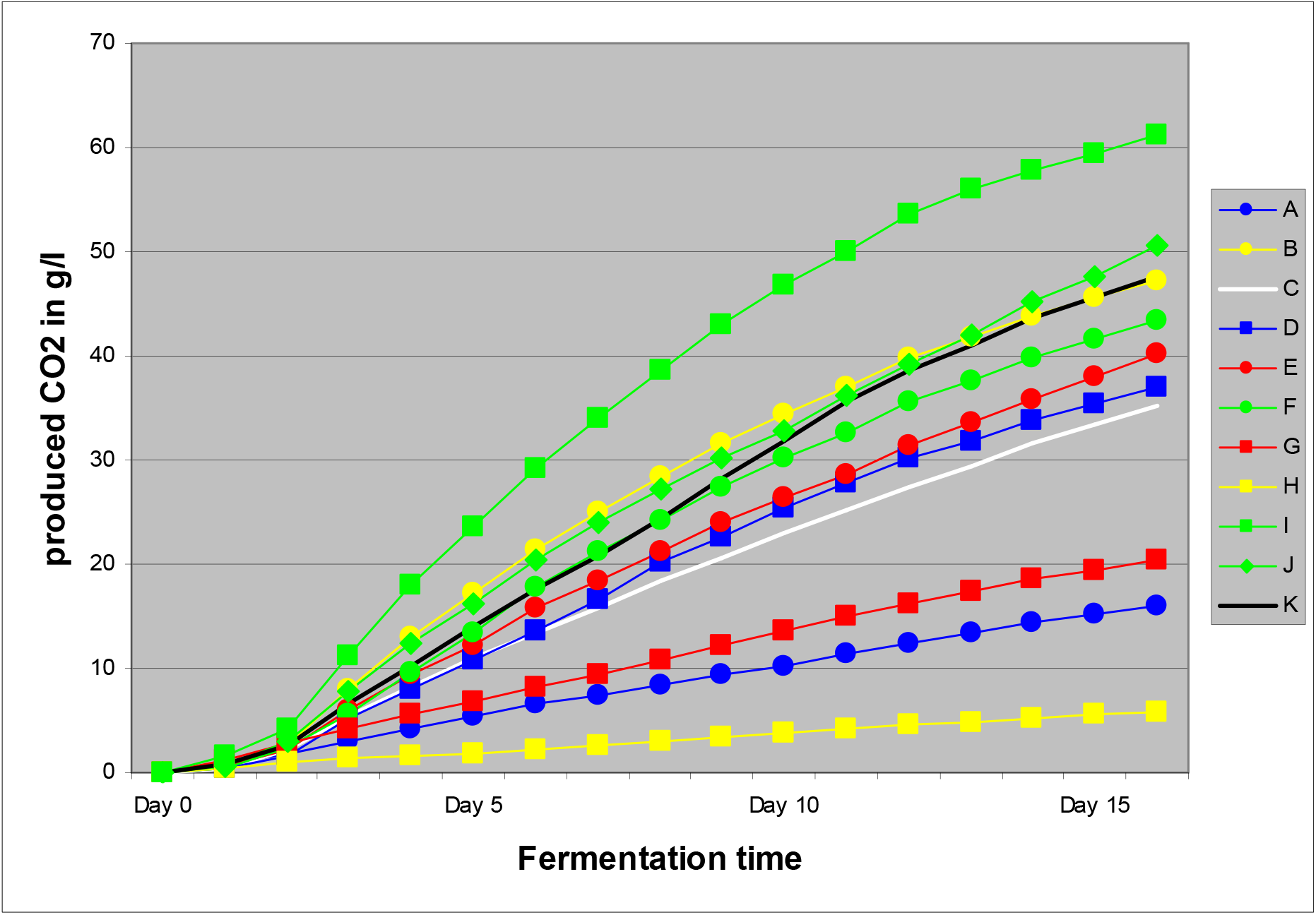
Comparison of fermentation efficiency by visualising the amount of carbon dioxide produced during fermentation. Red: vinified with yeast isolated from Speckbirne; yellow: isolated from Dorschbirne; blue: isolated from Wasserbirne; green: from Grüner Pichlbirne; white: from apple juice; black: from grape juice.

Principal component analysis (PCA), a multivariate method, was used to analyse which fruit wines differ significantly in terms of total basic chemistry (Fig. 2). In agreement with the previous analyses, it is not the yeasts obtained from grape or apple juice that differ the most. Most yeasts form a loose cluster, but four fruit wines do not belong to this cluster.

**Fig. 2:**
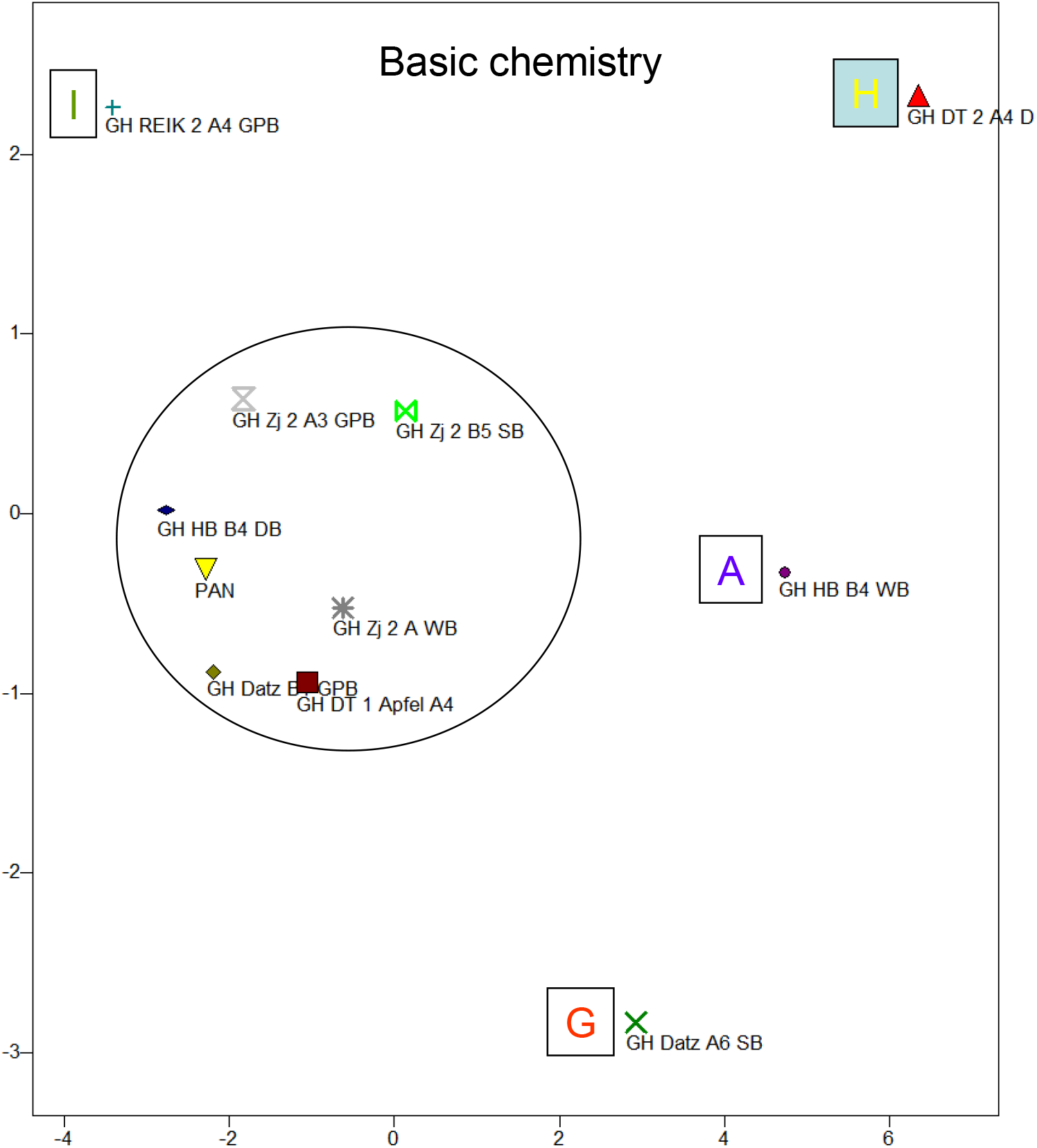
Principal component analysis of the basic composition of fruit wines produced with different yeasts.

GH DT 2 A4 DB wine (**H**), fermented using a yeast selected from fermenting Dorschbirne pear juice, has the highest density and highest glucose and fructose contents, as well as the lowest alcohol concentration. The glycerine content is also the lowest. There was practically no fermentation activity, but the yeast-available nitrogen was fully utilised.

GH HB B4 WB fruit wine (**A**) has a particularly high sugar-free extract, but very little total, titratable acid, as well as citric and malic acid. The yeast-available nitrogen is exceptionally high, the metabolic efficiency is modest.

GH Datz A6 SB fruit wine (**G**) with a yeast isolated from Speckbirne juice is characterised by modest metabolic efficiency with low nitrogen consumption. The values for sugar-free extract and also for volatile acid are comparatively very high. The pH-value is relatively low.

These (**A, G, H**) are the three yeast variants that were excluded from the experiment.

The GH REIK 2 A4 GPB pear wine (**I**), fermented with a yeast obtained from fermenting Grüne Pichlbirne fruit juice, has the lowest density and practically no sugar. There is no volatile acid and the pH value is higher than in all other test wines. However, the citric acid concentration is still the highest. In line with the excellent metabolic efficiency, no more yeast-available nitrogen can be detected. Yeast **I** is the most efficient fermenting yeast.

In general, the alcohol content of the pear wines is quite low at an average of 4% (Table 3), but this is due to the starting substrate. The comparative yeast Pannonia, isolated from fermenting grape juice, also only reached 4.9%.

The wines also do not cluster according to the pear variety from which the yeast was isolated (Fig. 3).

**Fig. 3:**
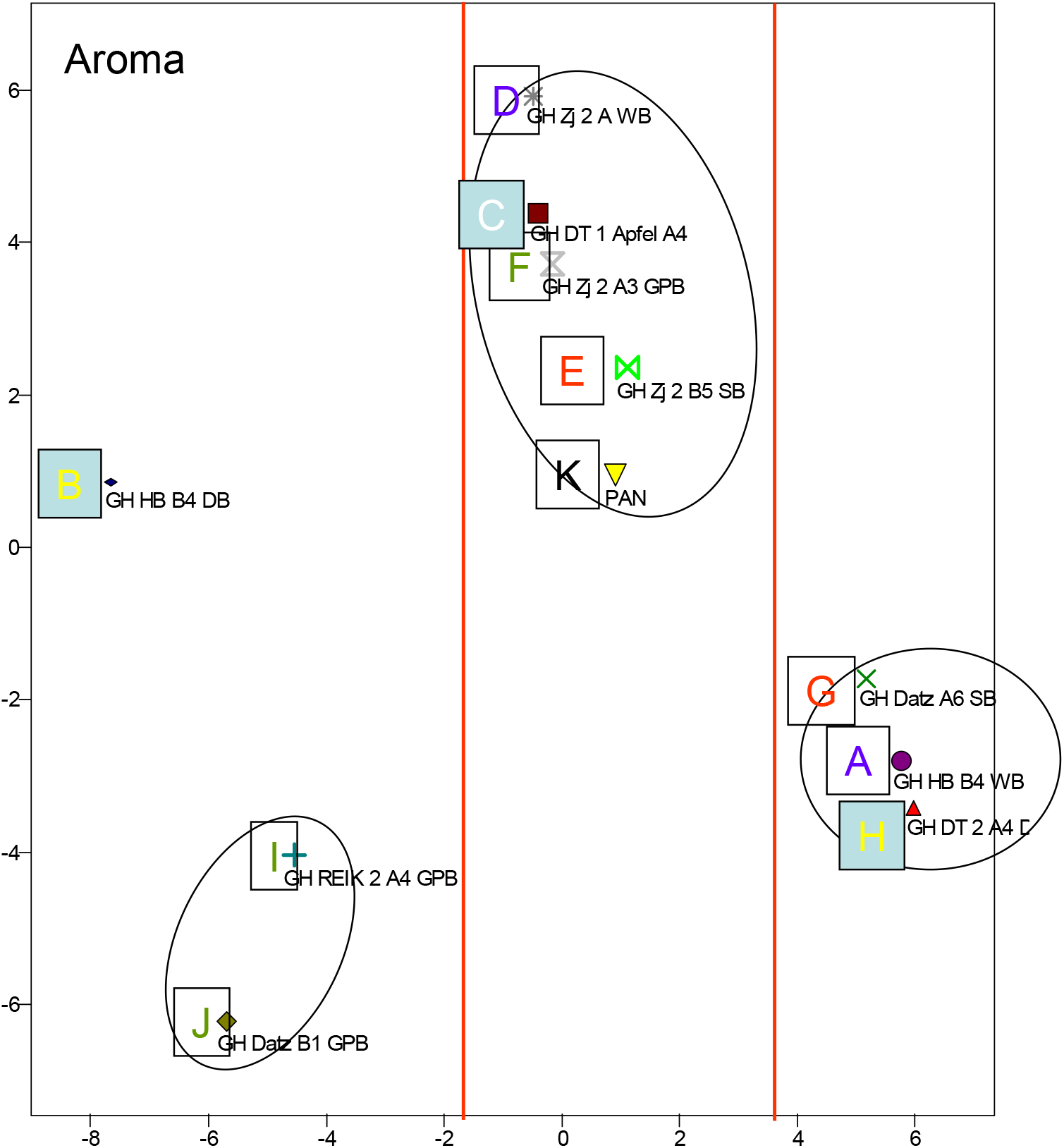
Principal component analysis (PCA) of the aroma spectra of pear wines produced with different yeasts (colour of capital letters: yellow: isolated from Dorschbirne pear; blue: isolated from Wasserbirne; green: from Grüner Pichlbirne; white: from apple juice; black: from grape juice).

In terms of flavour, four separate clusters are formed. At first glance, it looks as if basic chemistry and flavour spectrum have nothing in common, but the arrangement of the fruit wines along the first principle axis (abscissa) of Fig. 3 is strongly influenced by the metabolic efficiency of the fermenting yeasts. The three fruit wines on the left in Fig. 2 have a relatively high alcohol content on average (5.5%) and a correspondingly low sugar concentration (glucose: 0.8g/l; fructose: 10.3g/l). The fruit wines in the middle section of Fig. 3 also lie between the others in terms of these values (alcohol: 4.6%; glucose: 3.1g/l; fructose: 26g/l), while the three fruit wines shown on the right are characterised by a low ethanol content (1.3%) and high sugar concentration (glucose: 13.9g/l; fructose: 55.5g/l).

Fig. 4 shows an aroma diagram of the fruit wines (for a more detailed explanation, see the corresponding legend). The fruit wines whose yeasts have shown a high metabolic activity are mainly characterised by high concentrations of some ethyl esters and other esters (**B, I, J** in Fig. 3). The yeast **B** fruit wine, which is shown separately from the other two in PCA Fig. 3, also has relatively high concentrations of group 1 esters, which links it to the wines fermented with yeasts that showed average metabolic activity. However, these are also characterised by relatively high concentrations of octanoic and decanoic acid, while some of the ethyl esters are only present in small quantities. The group of fruit wines fermented by metabolically slow yeasts (**A, G, H**) generally have low concentrations of aroma compounds, although some of them may well be present in larger quantities, e.g. acetic acid ethyl ester or hexanol.

**Fig. 4:**
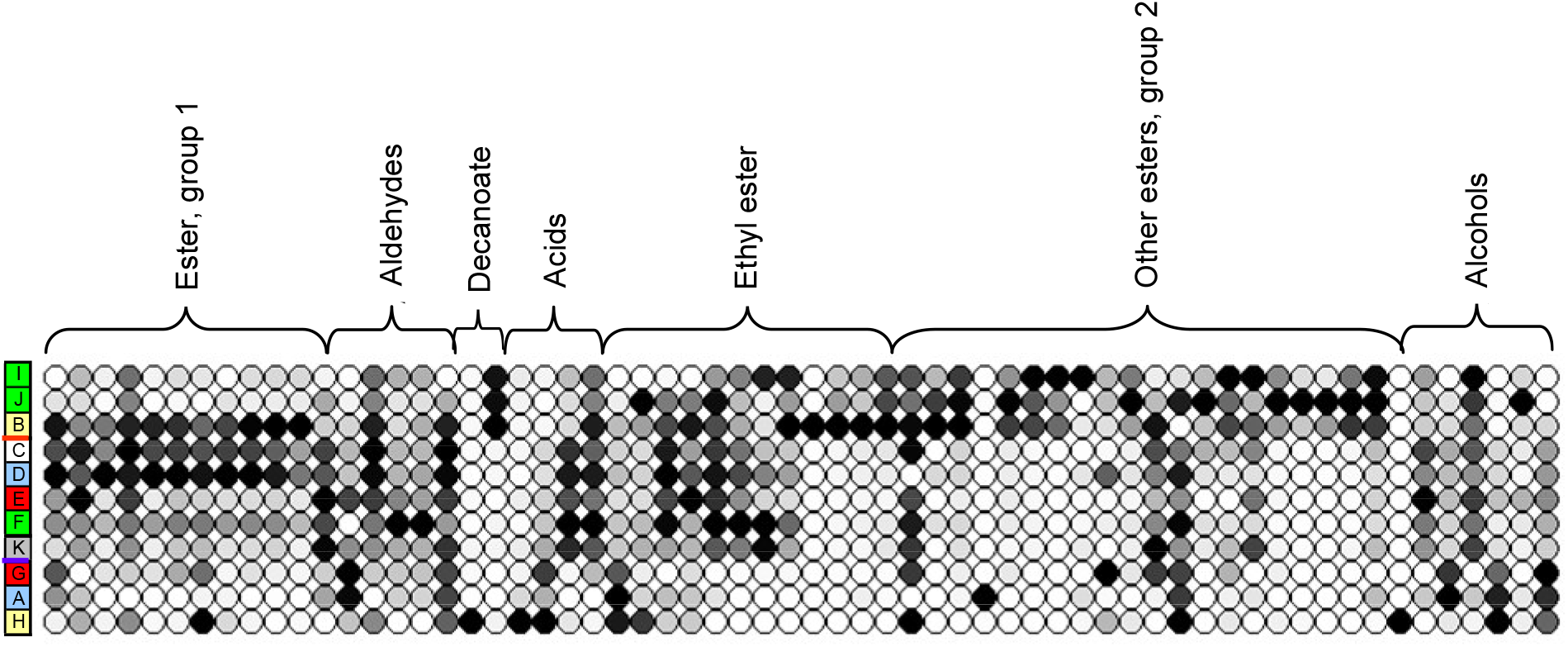
Aroma diagram of the fruit wines analysed, sorted according to the main component analysis (Fig. 3). The aroma substances are also sorted according to chemical groups. Within a column, all values are minimum-maximum scaled, i.e. the lowest value is represented by a white circle, the highest by a black one. All others are shaded grey. The aroma diagram provides no information about the absolute values, nor about the olfactory significance of the volatile substances.

### Fermentation in a 100-litre tank without comparative yeasts

Only the six best pear yeasts were used and no comparative yeasts. The fermentation process is characterised by a metabolically induced, steep rise in temperature at the beginning; some yeasts reach a temperature increase of 4°C. Yeast **B**, isolated from fermenting Dorschbirne pear juice, shows a particularly active metabolism and thus likely a corresponding fermentation rate, **E**, isolated from a Speckbirne pear fermentation, a particularly low one. This agrees with the observations from microvinification, if the elimination of three yeasts that had an even lower metabolic rate is taken into account.

The change in sugar gradation also shows this correlation (Fig. 6). Sugar degradation is particularly rapid with **B** and very slow with **E**.

**Fig. 5:**
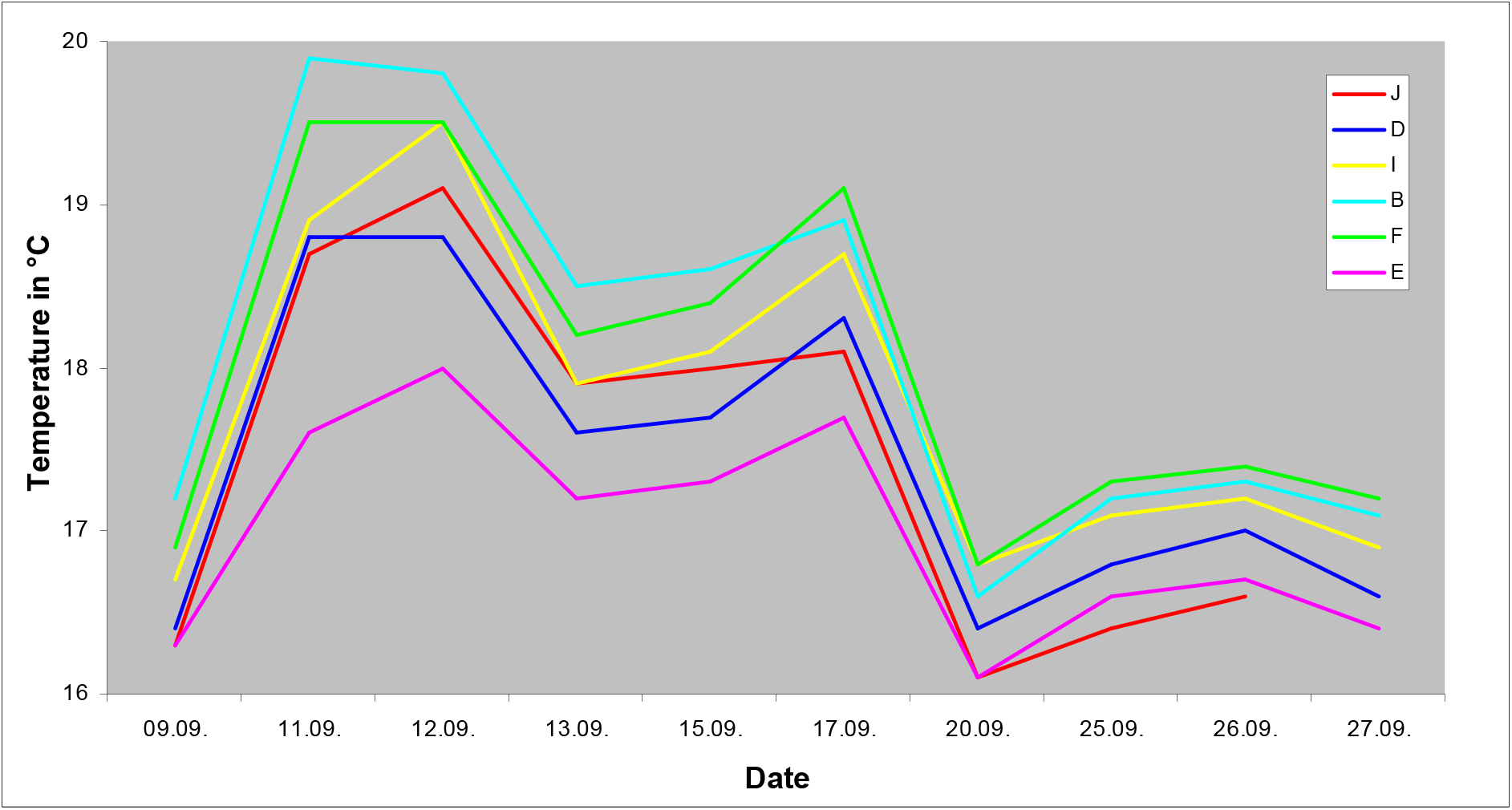
Temperature curves during vinification.

**Fig. 6:**
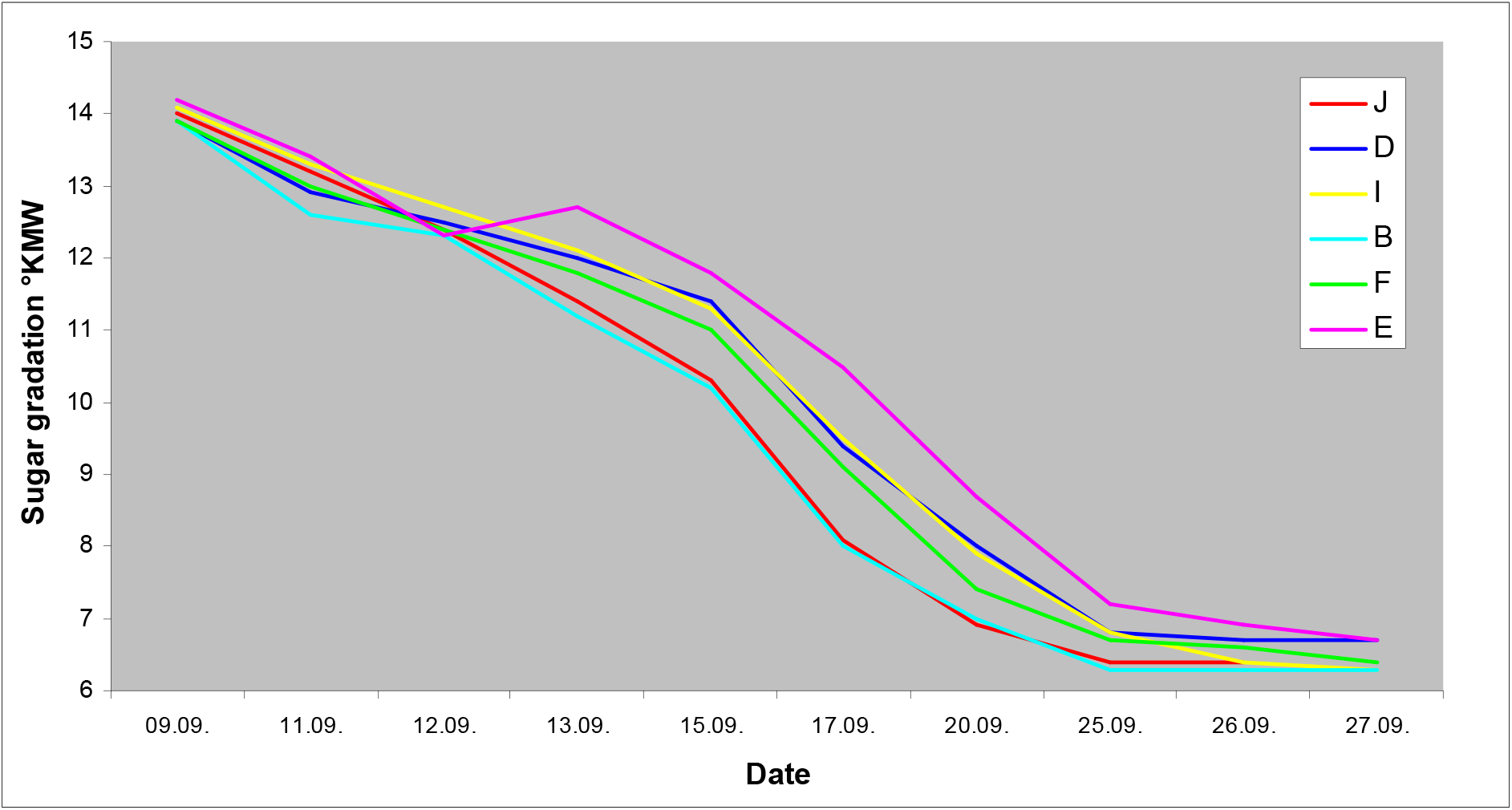
Change in sugar gradation during vinification.

As the basic analysis shows, **B, I** and **J**, the latter two isolated from fermenting Grüne Pichelbirne juice, ultimately metabolised all of the sugar, which is due more to persistent than rapid fermentation, particularly in the case of **I**. Glucose was completely metabolised by all yeasts, but a maximum of 1.7 g/l of fructose (**F** isolated from Grüne Pichlbirne) of the original 99.8 g/l was still present. In the fermentation tanks, as can be concluded from the high pH-values, acid degradation took place in **E** ("Speckbirne", pH 3.89) and **D** ("Wasserbirne", pH 3.82), whereas in **I** and **J** there was hardly any reduction in the acid concentration compared to the original product. The difference is particularly pronounced for the total titratable acid, but hardly present for the volatile acid. The concentration of malic acid is relatively high in **I** (3.9 g/l) and **J** (4.1 g/l) and particularly low in **E** (0.4 g/l). As expected, tartaric acid could not be detected. The citric acid concentration in all wines was in the range of 1.3 g/l to 1.4 g/l. The highest glycerol production was observed with the relatively fast fermenting yeast **J** (7g/l), followed by **B** (6.3 g/l), while **D** (Wasserbirne) performed worst with 4.5 g/l. The yeast-available nitrogen was completely consumed by **J**, while **B** and **D** were more economical here (11 mg/l and 10 mg/l respectively from the original 51 mg/l). The wine with the lowest glycerol concentration, **D** wine (produced with yeast from Wasserbirne juice), was rated the worst by the tasting committee (3:3 acceptance to rejection). Where a ranking was made, **B** wine was unanimously rated best and **D** worst. All other wines were rated comparably well.

The chemical analysis of the aroma compounds of the fruit wine (acetates, ethyl esters, higher esters, higher alcohols, free carboxylic acids, ketones) was carried out using gas chromatography/mass spectrometry (GC/MS). The primary evaluation consisted of a comparison of the aroma spectrum of the fruit wines with that of a white wine (Weißer Storch, Lenz Moser), which is described as harmonious and neutral in odour (Fig. 7). Compared to the white wine, the fruit wines are dominated or even exclusively characterised by aromas that are described as musty, sweet, floral and fruity (like honey), whereas the ethereal, fruity and green tones are more prominent in the white wine.

**Fig. 7:**
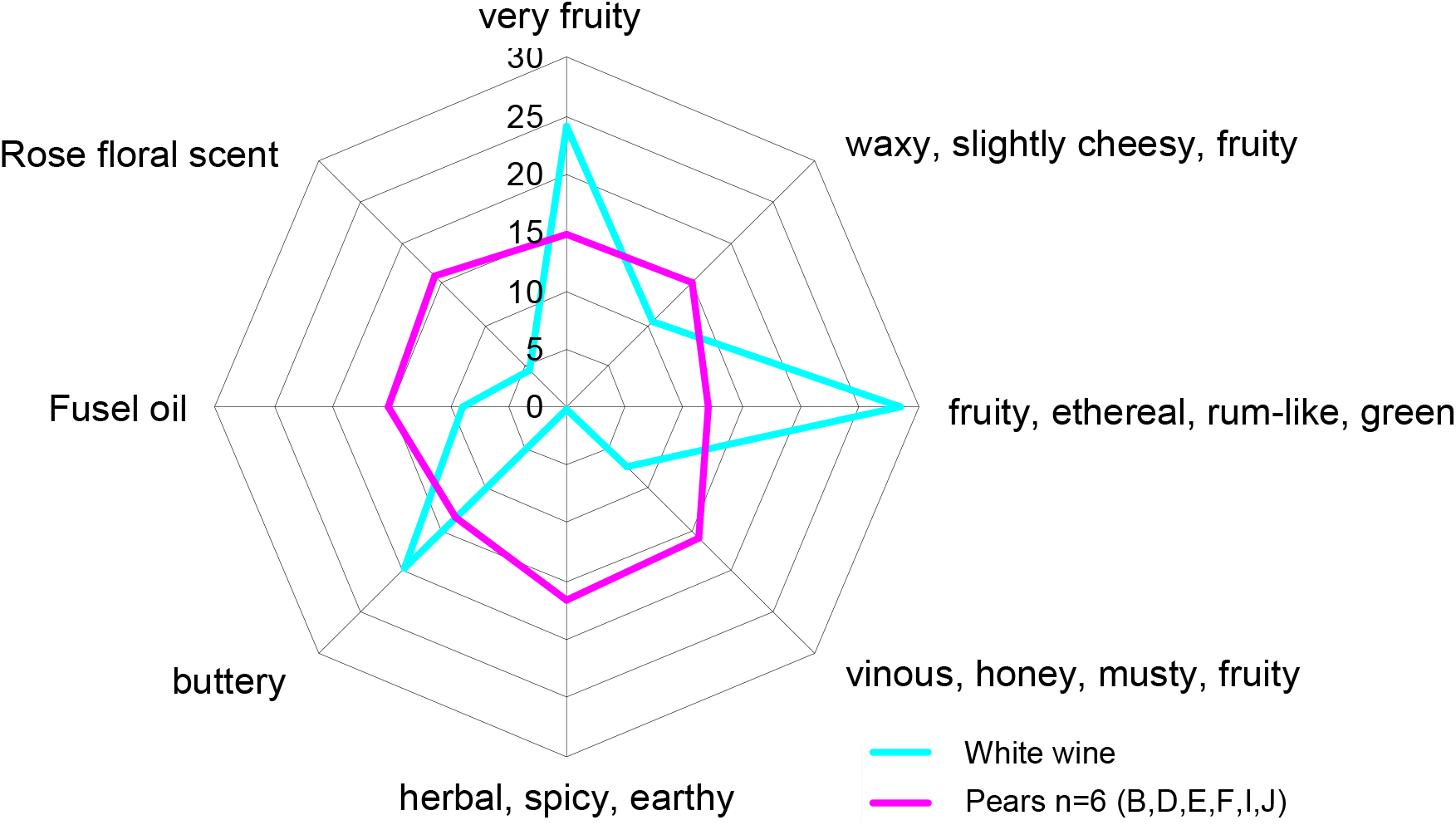
Comparison of the flavours of white wine and pear fruit wine.

If the aroma compounds examined are grouped according to the odor perception triggered by the individual components, characteristic differences emerge when comparing the fruit wines. For example, the aroma components of the fruit wine that was fermented with **B** yeast (isolated from Dorschbirne juice), which received a very good rating, are extremely well balanced (Fig. 8). In comparison, the worst-rated fruit wine (**D**, isolated from Wasserbirne juice) had a particularly strong planty, spicy, earthy note, which was partly due to the presence of 3-octanone, the smell of which was described as planty, sweetish and mushroomy. The highest concentration of butyric acid of the six fruit wines caused a further negative aroma perception. These aromas have a negative influence on the perception of fruit wines.

**Fig. 8:**
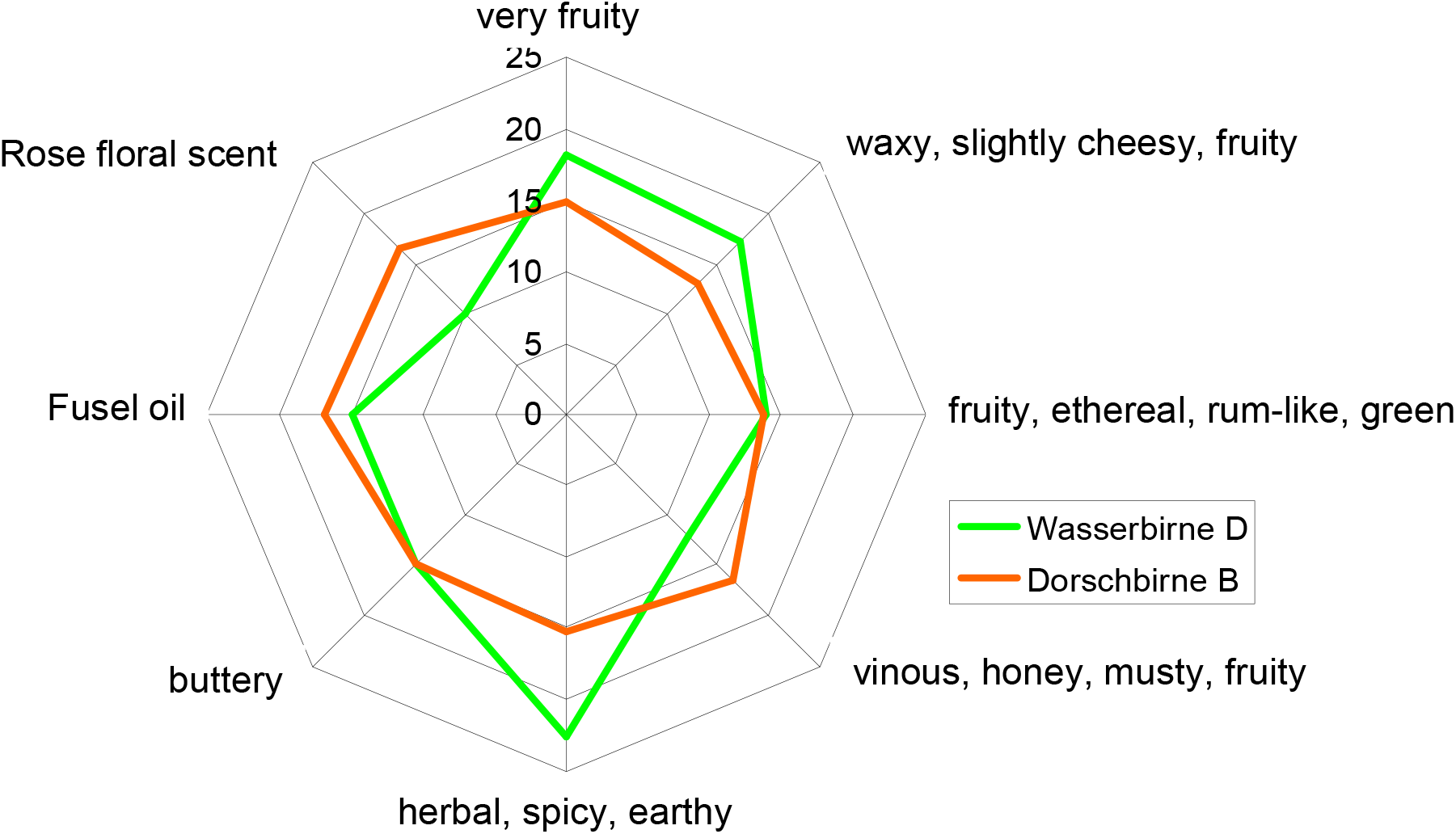
Comparison of the aromas of pear wine fermented with yeasts B (isolated from Dorschbirne juice) and D (isolated from Wasserbirne juice).

The aroma diagrams of all other pear fruit wines are summarized in Fig. 9.

**Fig. 9:**
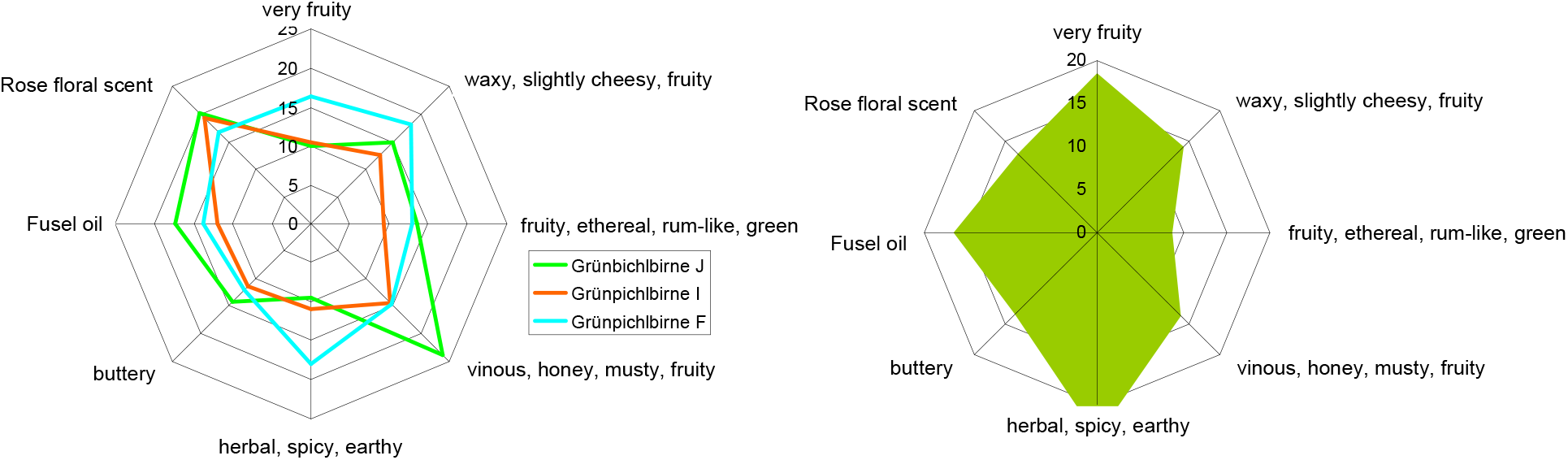
Aroma diagram of the pear fruit wines not discussed in detail in the text (left: Fermentation using yeasts isolated from Grüne Pichelbirne, right: from Speckbirne).

### Fermentation in 25-liter glass flasks and 100-liter tanks with comparative yeasts

After the pre-selection, one yeast each was left over, which had been isolated from Wasserbirne (**D**) and Speckbirne (**E**) (300ml microvinification); of those isolated from Dorschbirne, yeast **B** proved to be particularly active during vinification in the 100-liter tank. It was further cultivated into two strains ("Yeast_1" and "Yeast_2"). Of the yeasts isolated from Grüne Pichlbirne juice, **I** was selected for its metabolic efficiency in the 100-liter tank. In addition to the five native yeasts, four commercially available "grape-derived" pure yeasts were used: Actiflore Rose, Filtraferm Tropic, Oenoferm Freddo and Oenoferm X-Thiol. Fermentation took place for all nine yeasts with simple repetition in 25-liter glass flasks (18 wines) at the Haidegg experimental station, again in the following year; in addition, the "new" yeasts 1 and 2 were used for vinification in 100-liter tanks (in the Mostviertel region).

#### Tasting

To present the tasting results, the number of votes in favor of the marketability of the wine tested was divided by the size of the tasting panel (seven people), multiplied by two and the value one subtracted, resulting in an assessment number between one (all tasters consider the wine assessed to be marketable) and minus one (no one considers it to be marketable). Fig. 10 shows the result; the individual complaints are discussed below.

**Fig. 10:**
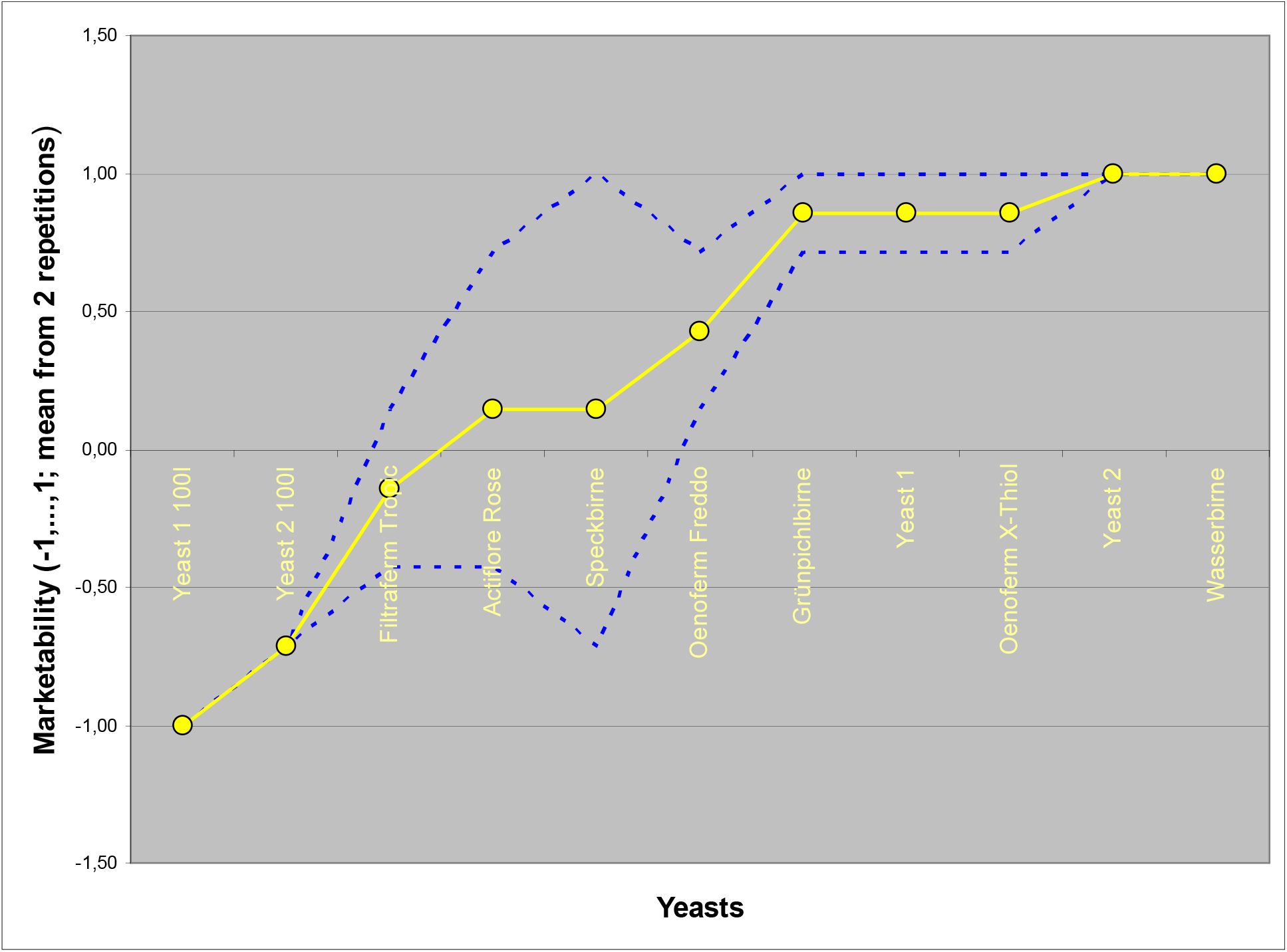
Result of the pear wine tasting. The yeasts used to ferment the pear juice are partly commercially available pure yeasts for grape wine production and partly new varieties developed within the project. Yellow: average of two replicates, blue: single wine.

The wines produced in 100-liter tanks received the worst ratings. The pear juice fermented with yeast_1 was assigned the off-flavours "volatile", "bumpy" and "impure", as was the wine produced with yeast_2, which was also described as "burnt". The fact that these odour and taste defects cannot be attributed to the yeast used is shown by the wines produced with the same yeasts in the glass flask. Yeast_1 was criticized once (out of 14), yeast 2 not at all. There was also no complaint for the yeast isolated from Wasserbirne juice or its fermentation product and only one complaint ("unclean") for "Grünpichlbirne". The two wines produced with Speckbirne yeast were judged very differently, one as completely faultless, while the other was found to have faults, namely "fermented" and "bockser". This means that fermentation was delayed and this wine is the only one to be criticized for the error "not yet fully fermented". Apart from this, it can be stated that in the Haidegg experiment all yeasts isolated from fermenting pear juice and cultivated purely were considered very suitable for pear wine production. Of the commercially available yeasts isolated from fermenting grape juice, however, this only applies to Oenoferm X-Thiol, a hybrid pure yeast for which the manufacturer claims the ability to increase the fruity thiol aroma as a characteristic feature. Only once is "styrene" (= smell of plastic) listed as a fault.

All other yeasts intended for grape wine production performed worse in the tasting than those isolated from fermenting pear juice. Filtraferm Tropic, a *Saccharomyces cerevisiae* yeast that is supposed to release exotic fruit aromas, appears particularly unsuitable, but is criticized under the test conditions due to defects in the wines produced such as "styrene", "medicine" and "reductive". Actiflore Rose (according to the manufacturer "for grape varieties with low aroma potential") was also judged only slightly better ("styrene", "medicine", "reductive", "foreign tone"). The wines produced with Oenoferm Freddo (*Saccharomyces cerevisiae* var. bayanus), a yeast with a very high fermentation strength, were criticized for "medicine", "reductive" and "styrene" off-flavours, but not rejected. All in all, it can be said that the yeasts isolated from fermenting pear juice and cultivated purely cause fewer faults in the fermentation of pear juice than the commercially available yeasts tested here. In the following, we will examine whether there is a fundamental difference between pure yeasts for grape wine production and yeasts isolated from fermenting pear juice.

### Basic chemistry

In the following Fig. 11, the relative concentrations or values are shown, a white circle therefore does not symbolize the concentration zero but the minimum concentration in the wine ensemble, a black one the maximum. The values of a column are therefore minimum-maximum scaled. The yeasts are sorted according to their affiliation to their selection medium ("grape yeast": fermenting grape juice; "pear yeast": fermenting pear juice).

**Fig. 11:**
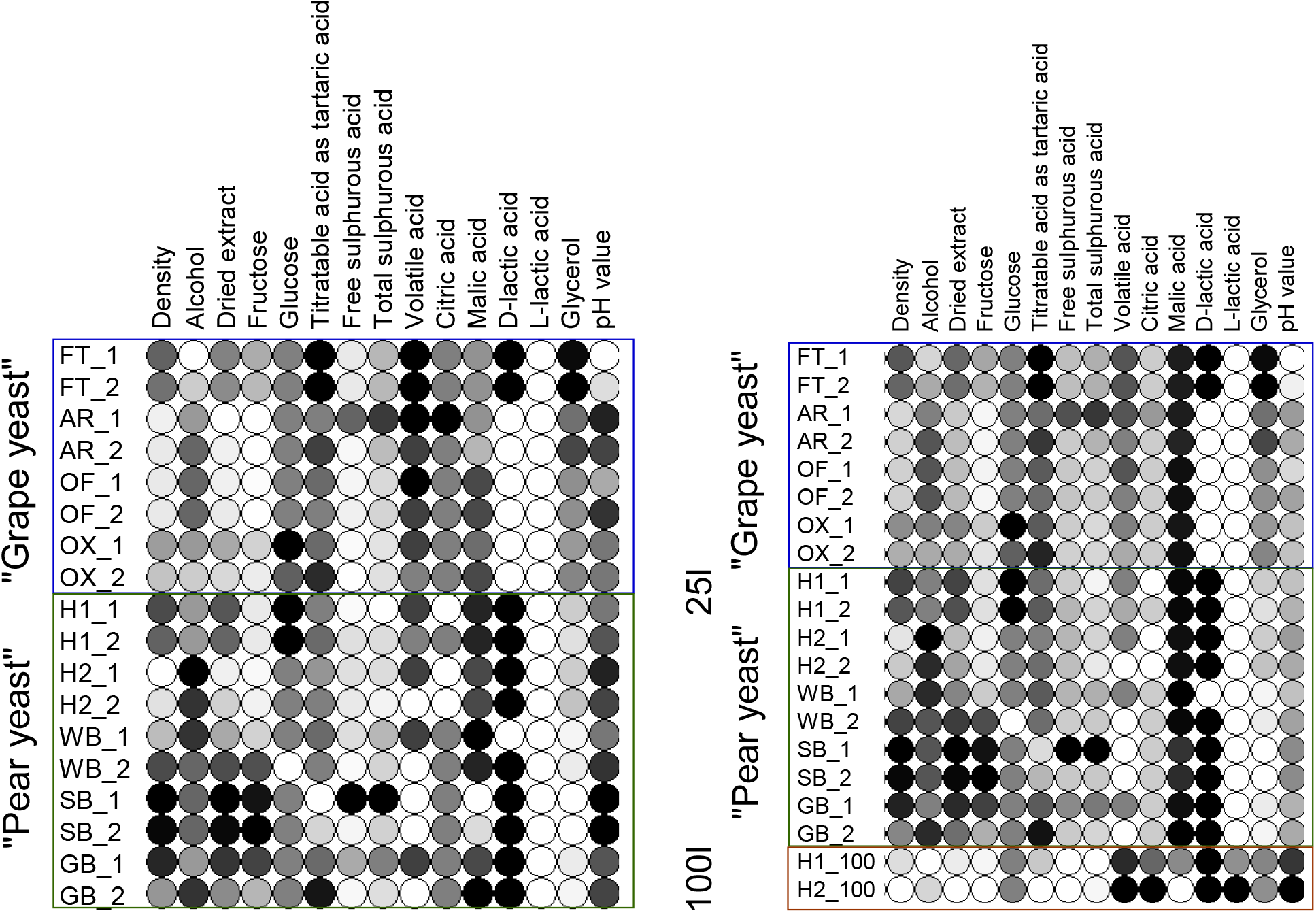
Basic chemistry of the test wines produced from the same pear juice with different yeasts. Left: Wines that were fermented in 25-liter containers. Right: Additionally those produced in 100-liter tanks. The abbreviations mean: AR = Actiflore Rose, FT = Filtraferm Tropic, OF = Oenoferm Freddo and OX = Oenoferm X-Thiol, H1 = Yeast 1 (Dorschbirne), H2 = Yeast 2 (Dorschbirne), WB = Wasserbirne, SB = Speckbirne, GB = Grünpichlbirne.

The wines undoubtedly have a pronounced individuality; in some cases, even those produced as a repetition under identical test conditions (including the same yeast) differ. Nevertheless, the majority of differences in basic chemistry can be identified between the wines of the different yeast selection media grape and pear juice. As Fig. 11 on the left shows, the density and dry extract content, as well as the fructose concentration, are generally higher in some "pear yeast" wines, i.e. these yeasts were less fructophilic than the "grape yeasts". Note, however, that the fructose content in the initial substrate is considerably higher than the glucose concentration (Table 1). The glucose consumption, on the other hand, was comparable in both groups, only slightly reduced in yeast 1. The production of malic acid is forced in the "pear yeasts": the wines have a content of 3.8g/l - 4g/l, while the values in the "grape yeast" wines are between 3.5g/l and 3.8g/l. The D-lactic acid concentrations are also slightly higher in the majority of "pear yeast" wines, at 0.1g/l, while they are not detectable in most "grape yeast" wines. Volatile acid (0g/l to 0.4g/l), glycerol (3.4g/l to 5.9g/l) and YAN (0mg/l to 15mg/l), on the other hand, are present in higher quantities in most "grape yeast" wines.

Fig. 11, right, also includes the wines produced in the 100-liter tank. They differ in terms of lower density, alcohol content, but also sugar concentrations and malic acid, as well as increased values for volatile and citric acid, L-lactic acid and pH-value.

The extent to which the wines are similar (or different) in terms of their basic data was investigated using a multivariate statistical method, the principal component analysis (Fig. 12). Although the individual wines differ quite clearly from each other, often even the two produced with the same yeast, the second main axis in particular separates the two groups "grape yeast" and "pear yeast" surprisingly clearly. The (more important) first main axis, on the other hand, primarily separates wines produced in 25-liter glass flasks from those produced in 100-liter tanks.

**Fig. 12:**
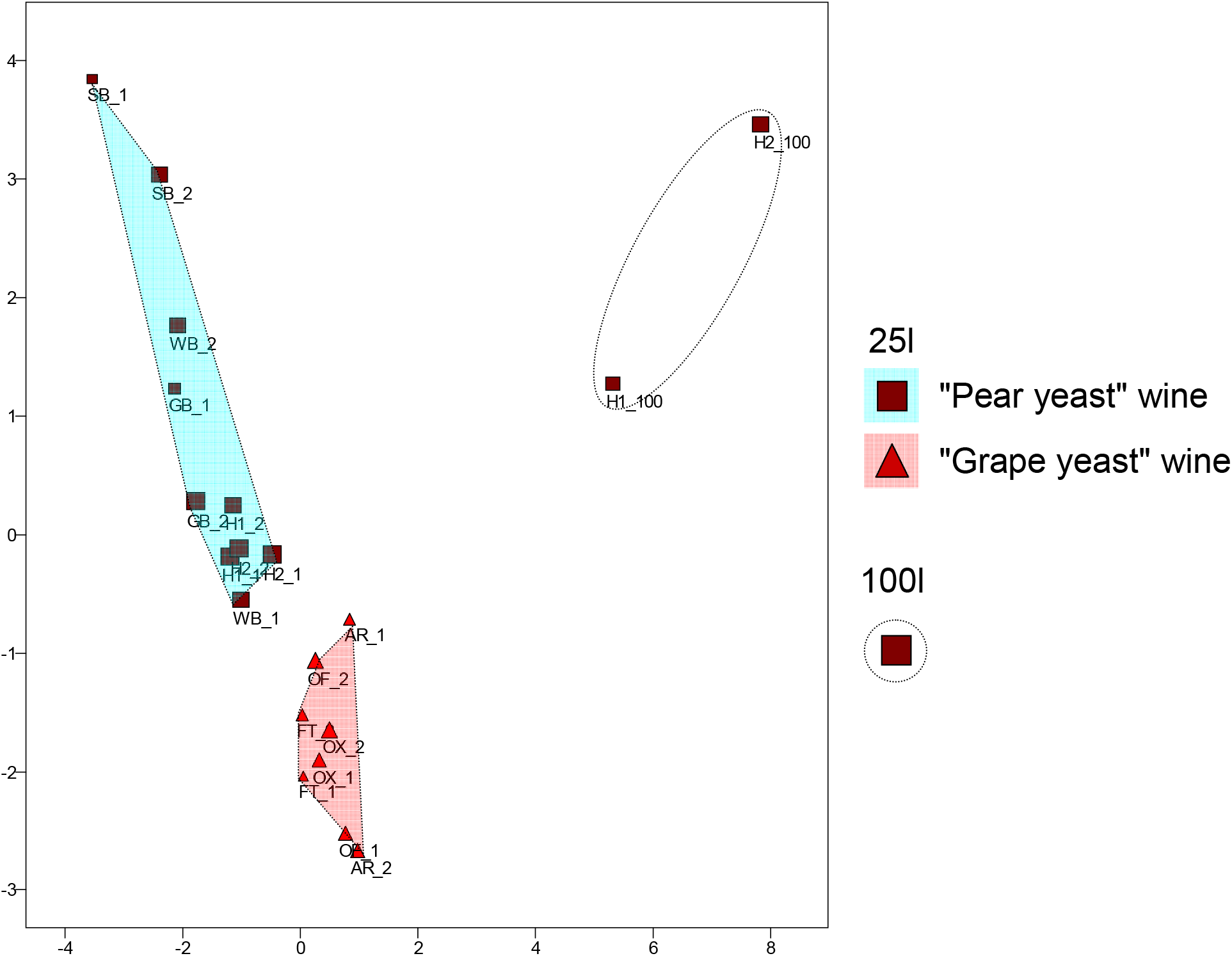
Principal component analysis of the basic chemical data of wines produced with different yeasts isolated from fermenting grape juice on the one hand and fermenting pear juice on the other.

### Aroma spectrum of the wines

39 aroma components typical of pear wines were selected for this study. Table 4 lists the description of the aroma perception that they trigger in most people in isolation. They are divided into aroma groups according to this perception.

**Table 4:** Aroma substances and their flavour description as isolated components.

| Substance | Aroma group | Aroma description (The good scents company) |
| --- | --- | --- |
| 1 iso-amyl acetate | Pear aroma | sweet fruity banana solvent |
| 2 n-pentyl acetate | Pear aroma | ethereal fruity banana pear banana apple |
| 3 n-hexyl acetate | Pear aroma | fruity green apple banana sweet |
| 4 Ethyl 2,4-decadienoate | Pear aroma | green waxy pear apple sweet fruity tropical |
| 5 2-phenylethyl acetate | flowery, rose | floral rose sweet honey fruity tropical |
| 6 2-phenylethanol | flowery, rose | floral rose dried rose flower rose water |
| 7 ethyl lactate | buttery | sharp tart fruity buttery butterscotch |
| 8 Acetoin | buttery | sweet buttery creamy dairy milky fatty |
| 9 2-ethylhexanol | citrus | citrus fresh floral oily sweet |
| 10 Limonene | citrus | citrus orange fresh sweet |
| 11 n-propyl acetate | fruity | solvent celery fruity fusel raspberry pear |
| 12 butyl acetate | fruity | ethereal solvent fruity banana |
| 13 heptyl acetate | fruity | fresh green rum ripe fruit pear apricot woody |
| 14 ethyl acetate | fruity | ethereal fruity sweet weedy green |
| 15 ethyl propionate | fruity | sweet fruity rum juicy fruit grape pineapple |
| 16 ethyl 2-butenolate | fruity | pungent chemical diffusive sweet alliacious caramel rum |
| 17 ethyl isobutyrate | fruity | sweet ethereal fruity alcoholic fusel rummy |
| 18 ethyl butyrate | fruity | fruity juicy fruit pineapple cognac |
| 19 ethyl hexanoate | fruity | sweet fruity pineapple waxy green banana |
| 20 ethyl heptanoate | fruity | fruity pineapple cognac rum wine |
| 21 ethyl octanoate | fruity | fruity wine waxy sweet apricot banana brandy pear |
| 22 ethyl nonanoate | fruity | fruity rose waxy rum wine natural tropical |
| 23 ethyl 9-decenoate | fruity, sweet | fruity fatty |
| 24 ethyl decanoate | fruity, sweet | sweet waxy fruity apple grape oily brandy |
| 25 ethyl dodecanoate | fruity, sweet | sweet waxy floral soapy clean |
| 26 ethyl tetradecanoate | fruity, sweet | sweet waxy violet orris |
| 27 ethyl hexadecanoate | fruity, sweet | mild waxy fruity creamy milky balsam |
| 28 propyl hexanoate | fruity, sweet | sweet fruity juicy pineapple green tropical |
| 29 iso-amyl alcohol | Fusel oil | fusel oil alcoholic whiskey fruity banana |
| 30 hexanol | green | ethereal fusel oil fruity alcoholic sweet green |
| 31 heptanol | green | Musty leafy violet herbal green sweet woody peony, weak alcohol |
| 32 n-octanol | green | waxy green orange aldehydic rose mushroom |
| 33 iso-octyl acetate | Herbs, forest | earthy herbal humus undergrowth |
| 34 n-octyl acetate | Herbs, forest | green earthy mushroom herbal waxy |
| 35 3-octanone | Herbs, forest | fresh herbal lavender sweet mushroom |
| 36 α-Terpinolene | Herbs, forest | fresh woody sweet pine citrus |
| 37 Acetic acid | sour, pungent | sharp pungent sour vinegar |
| 38 Styrene | Pollution |  |
| 39 Xylene | Pollution |  |

Only a few are rated as particularly unpleasant in isolation, e.g. acetic acid, styrene and xylene, ethyl lactate and acetoin. One of the aroma groups ("pear aroma") indicates which substances are particularly important for the pear flavour.

For the evaluation, a principal component analysis (PCA) was first carried out on the basis of the aroma components. The graphical representation (Fig. 13) of the position of the wines with regard to the first two principal components initially shows in the left-hand figure that the first principal component (PC1, horizontal axis) allows the wine aromas produced in the 100-liter tanks to be separated very well from those produced in the 25-liter containers (PC1 explains 33% of the variance). The second principal component (PC2, vertical axis), on the other hand, does not separate the aromas of "pear yeast" wines and "grape yeast" wines quite as clearly, but still quite well.

**Fig. 13:**
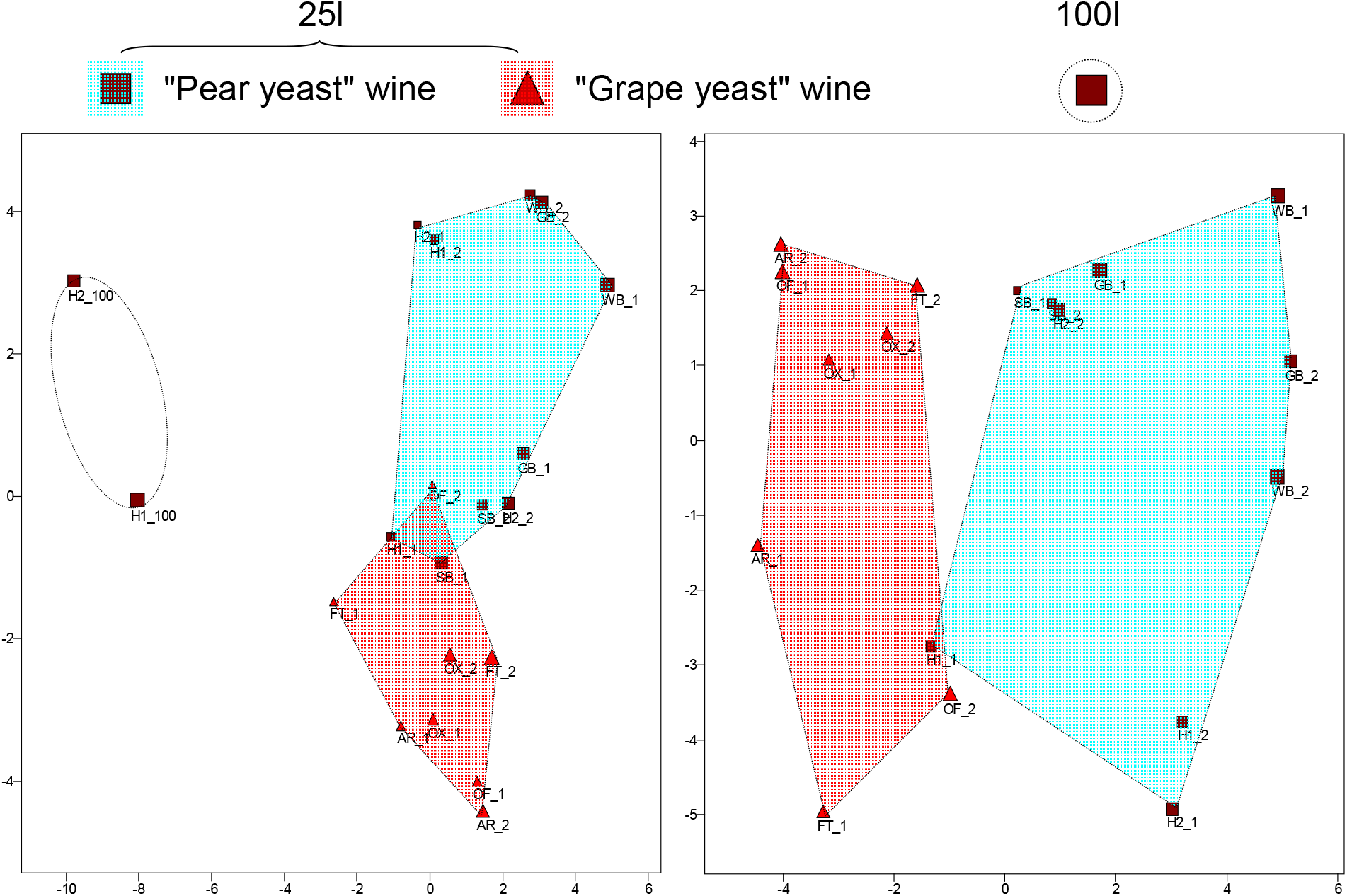
Principal component analysis based on the aroma components of wines produced with different yeasts isolated from fermenting grape juice on the one hand and fermenting pear juice on the other.

In the partial illustration on the right, PCA was carried out exclusively for the wines produced in the 25-liter glass flasks. Here it is PC1 that separates grape and pear yeast wines from each other, and does so better than in the left-hand partial figure (PC1, right, explains 29% of the total variance, PC2, left, only 21%). What has already been established for the basic chemistry, namely that the wines, often also the pairs, which were produced with the same yeast, appear quite different, also applies to their aromas and it is therefore all the more remarkable that the separation according to the aroma components is almost unambiguous. The pear wines are positioned along PC1 according to the equation:

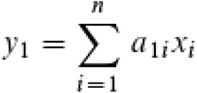

where x_i_ corresponds to the aroma concentration of the ith aroma substance. y is the value of the wine on the PC1 axis. The greater the amount of the constant a_1i_ calculated by the PCA, the more important the substance is for the positioning. Substances for which │a_1i_│>0.1 apply are discussed below. The sign provides additional information as to whether the concentration in the pear wine group is relatively high or low.

For the positioning of the wines along PC1 Fig. 13 left, some aroma components are particularly significant, i.e. for the separation between 100-liter tank wines and the 25-liter vinification. In addition to the main component analysis, this is also shown in the lower part of Fig. 14. According to this, fruity substances such as n_propyl acetate, butyl acetate, heptyl acetate, ethyl butyrate, ethyl hexanoate and ethyl octanoate, as well as fruity-sweet substances such as ethyl decanoate, ethyl dodecanoate and propyl hexanoate, and aromas from the "herbs, forest" group (n_octyl acetate, a_terpinolene) predominate in the 25-liter wines. In the 100-liter wines, substances from the "sour, pungent" (acetic acid), buttery (ethyl lactate, acetoin) and "green" (heptanol and octanol) aroma groups are more concentrated. However, there are also pleasant components that dominate, namely 2-phenylethanol ("floral, rose") and limonene ("citrus").

**Fig. 14:**
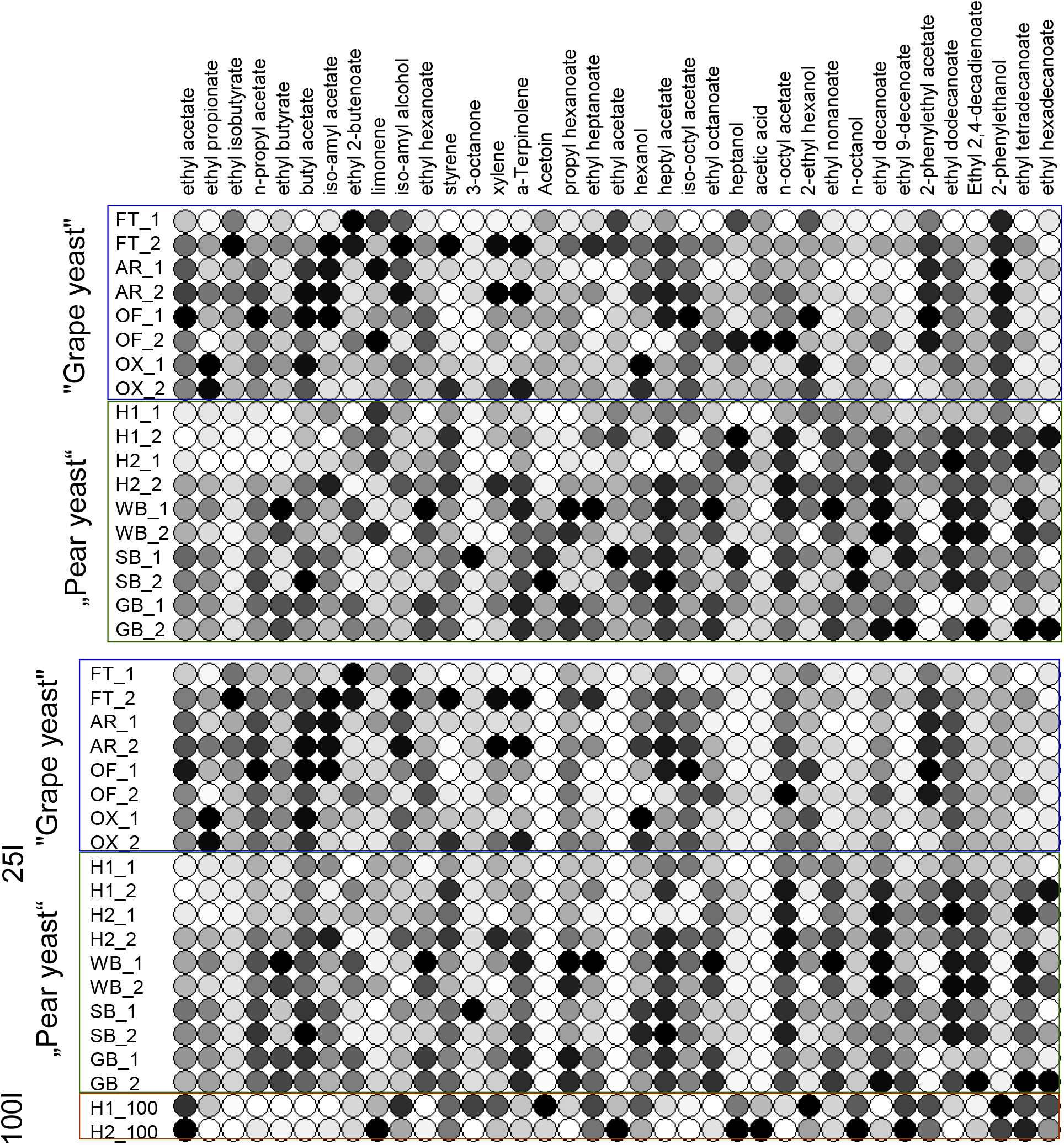
Aroma spectrum of the test wines produced from the same pear juice with different yeasts. Top: Wines fermented in 25-liter containers. Below: Additionally those that were produced in 100-liter tanks.

PC1 in Fig. 13, right, separates pear yeast wines from grape yeast wines. Some aromas from different groups are mainly responsible for the positioning along the first principal axis (see also Fig. 14, above). A higher concentration of the following substances is typical for grape yeast wines: From the "fruity" aroma group, these are butyl acetate, ethyl acetate and ethyl iso-butyrate; from the "pear aroma" group, iso-amyl acetate; from the "herbal, forest" group, iso-octyl acetate; from the "floral, rose" group, phenyl ethyl acetate and phenyl ethanol; from "sour, pungent", acetic acid; from "fusel oil", iso-amyl alcohol; and from the citrus group, ethyl hexanol.

In contrast, pear yeast wines are dominated by n-octyl acetate from the "herbs, forest" group, ethyl butyrate, ethyl hexanoate, ethyl heptanoate, ethyl octanoate, ethyl nonanoate from the "fruity" aroma group, ethyl 2-4-decadienoate from the pear group, ethyl 9-decenoate, ethyl decanoate, ethyl tetradecanoate, ethyl hexadecanoate, propyl hexanoate from "fruity, sweet", n-octanol from the "green" flavour group and styrene from "environmental pollution". The aroma of pear yeast wine is therefore characterized by significantly more fruity and fruity-sweet aromas. "Flowery-rose", on the other hand, tends to characterize grape yeast pear wines, and in both pear wine groups there are also substances that tend to cause negative olfactory or taste impressions in isolation.

